# Platelet Molecular Maturation Links Platelet Aging and the Platelet Storage Lesion

**DOI:** 10.64898/2026.02.03.703379

**Authors:** Cedric M.V. Bainton, Yale Santos, Fahima Mayer, Alexander Fields, Karl Zoghbi, Nasima Mayer, Danielle Williamson, Greg Chinn, Katelin C. Rahn, Javonn M. Musgrove, Kimberly A. Thomas, Jeff Sall, Susan M. Shea, Lucy Z. Kornblith, Roland J. Bainton

## Abstract

Human platelets change over their 7–10 day lifespan, yet the molecular mechanisms underlying platelet aging remain poorly defined. Using two independent RNA sequencing datasets of fluorescence-activated cell sorted young and old human platelets, we developed a unified transcriptomic model to characterize RNA metabolism across the platelet lifespan, which we termed platelet molecular maturation. This was applied to RNA sequencing data from room-temperature stored platelets (up to 7 days) and cold-stored platelets (7, 14, or 21 days). We identified highly concordant aging signatures, including 6,015 shared expressed genes and 2,008 shared differentially expressed genes (DEGs) with strongly correlated fold changes, demonstrating a conserved platelet aging program. Nucleotide-level analyses revealed preferential 3′-directed degradation among downregulated transcripts during endogenous platelet aging and room-temperature storage, supporting an organized RNA decay process that was correlated with platelet function changes. Room-temperature storage recapitulated platelet molecular maturation, showing concordance with aging-related gene expression changes and enrichment of downregulated gene sets. In contrast, cold-storage significantly attenuated platelet molecular maturation and 3′-directed degradation. A total of 669 genes were consistently differentially expressed between room-temperature and cold-stored platelets, while no DEGs were detected during cold-storage, indicating transcriptional stability. Platelet transcript stability in cold-storage correlated with preserved platelet hemostatic function. These findings establish platelet molecular maturation as a conserved, functionally relevant model linking endogenous platelet aging to platelet storage lesions and providing mechanistic insight into preserved platelet hemostatic function in cold-storage. This atlas of platelet RNA metabolism supports biomarker discovery and strategies to improve storage.

**KEY POINTS:**

- By integrating multiple high-quality RNA sequencing datasets with novel analytic approaches tailored to the biology of anucleate platelets, we show that platelet aging is not a passive process of transcript decay, but follows a structured and reproducible molecular trajectory both endogenously and in storage, which we term platelet molecular maturation.
- Storage temperature emerged as a dominant modifier of this trajectory, with cold-storage markedly slowing RNA metabolic kinetics and preserving transcripts associated with younger, more hemostatically competent platelets.
- Together, these findings provide mechanistic insight into the platelet storage lesion and identify transcriptomic features that may serve as biomarkers or therapeutic targets to extend platelet shelf life.

## INTRODUCTION

Human platelets have an average lifespan of 7-10 days, resulting in heterogeneous circulating populations that vary in size, phenotype, metabolic capacity, and, consequently, function.^1-10^ As platelets age, their metabolic capacity declines and apoptotic processes begin, increasing their potential for participation in coagulation, while impairing adhesion and aggregation behaviors.^11-13^ Although platelet phenotype and function change over the course of their lifespan, the molecular processes governing platelet aging remain poorly defined.^14^ Notably, young (reticulated, large) and old (non-reticulated, small) platelets exhibit distinct RNA profiles, with the young platelet transcriptome enriched for calcium signaling, aggregation responses to collagen, thrombin, and adenosine diphosphate, and α-granule mobilization compared to old platelets.^15,16^ Collectively, these studies suggest that the platelet transcriptome may play a central role in regulating endogenous platelet aging and governing platelet phenotype related to coagulation, inflammation, and immune regulatory functions.

Further complicating this issue is the impact of endogenous platelet aging on platelet quality and function under storage. Transfusion of stored platelets is an essential life-saving intervention for patients with a wide range of disease etiologies, accounting for approximately 2.2 million transfusions each year in the United States (US).^17,18^ However, a National Blood Collection and Utilization Survey found that 24% of manufactured platelet products (∼631,000) expired and had to be wasted before clinical use.^17,18^ Accordingly, efforts to advance our understanding of stored platelet phenotypes and function, as well as to optimize storage conditions, are of central importance.^17-19^ Platelet products are currently stored at room-temperature (5-7 days at 22°C), during which biochemical, structural, and functional deterioration occurs, referred to as the “platelet storage lesion”. The platelet storage lesion reduces post-transfusion survival, impairs clot formation, and limits usability to 5-7 days..^20^ Importantly, despite accumulating evidence that the platelet transcriptome plays a role in endogenous platelet aging and loss of function, the mechanisms by which the transcriptome changes and subsequently affects platelet function during storage are unknown.

Refrigeration, or cold-storage, of platelets (up to 14 days at 4°C)^21^ is one strategy to extend platelet shelf life and reduce wastage.^22-24^ An added benefit of cold-storage is that it preserves hemostatic function of platelets better than room-temperature storage, as assessed using laboratory techniques to measure platelet function.^25-28^ Cold-storage of platelets out to 14 days is used in a small number of centers under a US Food and Drug Administration (FDA) guidance,^21^ and clinical trials are underway to assess cold-stored platelet safety and efficacy for bleeding patients receiving platelet transfusions.^29-31^ However, the mechanisms regulating the preserved hemostatic function of cold-stored platelets remain incompletely defined. As a result, our capacity to strategically manipulate these mechanisms to prolong platelet storage and enhance transfusion outcomes remains fundamentally constrained. We have previously found that alterations in metabolic capacity may be associated with platelet functionality during storage^25^, suggesting that metabolic age could play a central role in the storage lesion. Interestingly, the enhanced hemostatic preservation capacity displayed by cold-stored platelets is surprisingly similar to the dichotomy of functions between young and old endogenous platelets. Limited investigations have assessed the transcriptome of room-temperature stored platelets,^32,33^ and none have examined the impact of cold-storage on the platelet transcriptome or its potential association with preserved platelet hemostatic function.

In this study, we integrated novel bioinformatic approaches and meta-analyses to develop a model of endogenous platelet aging driven by conserved platelet RNA metabolism, termed platelet molecular maturation. We then leveraged this to examine storage-associated changes in platelet RNA metabolism to demonstrate that cold-storage significantly preserves the platelet transcriptome and associated platelet function, compared to room-temperature storage.

## METHODS

All included human subjects research was approved by the University of Pittsburg Institutional Review Board (#21110093, respectively), with data and material sharing agreements between the University of Pittsburg and the University of California, San Francisco. Informed consent for platelet collection was obtained where applicable.

### Publicly Available RNA Sequencing Datasets for Development of the Platelet Molecular Maturation Model

We leveraged two high-quality split-sample publicly available RNA sequencing (RNAseq) datasets (European Nucleotide Archive PRJEB35689^2^ – referred to herein as first-senior author “Hille-Trenk”^15^ *[HT]* dataset; Sequence Read Archive GSE126448^1^– referred to herein as first-senior author “Bongiovanni-Bernlochner”^16^ *[BB]* dataset) of fluorescence-activated cell sorted (FACS) endogenous platelets at two opposing ends of the platelet lifespan (young versus old) to create our platelet molecular maturation model. Both HT and BB datasets used size and intensity of nucleic acid fluorescence to create gating windows for young (large) and old (small) platelets and were reanalyzed using our bioinformatic methods below.

### Collection, Functional Measurements, and Sequencing of Stored Platelets

Apheresis platelets (Trima Accel, Terumo, US; resuspended in plasma and ACD-A) were collected from healthy volunteer donors (N=5, University of Pittsburgh Institutional Review Board #21110093) in accordance with FDA regulations and Association for the Advancement of Blood and Biotherapies standards and guidelines. Units were split equally, with one part stored agitated at 22°C (room-temperature stored platelets) and the other unagitated at 4°C (cold-stored platelets) per current standard practice. Units were sampled at baseline (day 0) and 7 for room-temperature stored platelets and days 0, 7, 14, and 21 for cold-stored platelets. At these sampling timepoints, functional measurements and platelet RNAseq were performed as described below. Further details are available in the **Supplemental Materials**.

Complete blood counts and residual white blood cell counts were performed. Light transmission aggregometry (single and dual agonists; adenosine diphosphate (ADP), epinephrine, collagen, ADP+epinephrine, ADP+collagen) was performed (ChronoLog, US), and lag time, slope, maximum amplitude, and area under the curve recorded. Thrombin generation in response to 5pM tissue factor was measured via Calibrated Automated Thrombogram (Diagnostica Stago, US), and lag time, peak thrombin, and endogenous thrombin potential recorded. Viscoelastic testing was performed via EXTEM (extrinsic cascade) and FIBTEM (extrinsic cascade with platelet inhibition) tests (ROTEM delta, Werfen, US). Clotting time, clot formation time, maximum clot formation, and lysis index at 60 min were recorded. Further details are available in the **Supplemental Materials**.

Platelet isolation and RNAseq methods are detailed in **Supplemental Materials**. In brief, platelets were isolated from apheresis unit samples using negative selection (anti-CD45 magnetic beads), pelleted, resuspended in Trizol, and stored at -80°C. RNA was isolated from frozen samples via Qiamp RNA Blood Mini Kit (Qiagen, Germany), contaminating DNA removed with DNAse I, and sample quality assessed by Bioanalyzer (Agilent, US). Complementary DNA libraries were generated with 1-2 ng of RNA via the Ovation random primed isothermal amplification system (NuGen, US). Sequencing was performed at a depth of 100 to 400 million reads using a NovaSeq 6000 (Illumina, US). These RNAseq datasets are herein referred to as *room-temperature stored* and *cold-stored* datasets.

### Bioinformatic Methods

#### Sequence Alignment and Mapping

For all RNAseq datasets (*HT, BB, room-temperature stored*, and *cold-stored*), procedures are detailed in **Supplemental Materials**. In brief, quality was assessed with FastQC,^34^ read duplication counted using RSeQC,^35^ mapping classifications (coding, untranslated region [UTR], intronic, intergenic or ribosomal) measured using the Picard software suite,^36^ and all metrics visualized with MultiQC^37^. All datasets were aligned to the human genome (GRCh38 assembly) and mapped using STAR 2.6.0c^38^. Gene expression values were counted using featureCounts.^39^

#### Comparative Differential Gene Expression Analysis

For each RNAseq dataset comparison, gene expression changes were compared with edgeR^40^, with a |log2 fold change (FC)|>1 and false discovery rate (FDR) < 0.05 as thresholds for significance. Gene set enrichment analysis was performed with the *GSEA* module of the *GSEApy* Python library^41^. Further details are available in the **Supplemental Materials**.

#### RNA Metabolism Analysis

The anucleate nature and long half-life of platelets required new tools to examine RNA metabolism changes during platelet aging: specifically, the construction of a gene coverage map down to the nucleotide level and multiple data aggregation steps to elaborate the nature of RNA processing events. Therefore, we analyzed RNA metabolism by summation of the statistically differing individual nucleotides gained and lost over maturation (e.g., age and storage) which we have termed “Nucleotide Level Statistical Composite” or NLSC. The NLSC counts bulk nucleotide shifts that can be partitioned into negative or positive changed positions, which is useful for considering enzyme 5’ or 3’ location of nucleotide processing, and provides clues to the different types of RNA metabolism. Further details are available in the **Supplemental Materials**, including **Supplemental Figure 3**.

#### Integrated Data Analyses

Principal component analysis (PCA) was used to capture relationships between gene expression changes across genomic and functional measurement datasets. Gene expression was z-score normalized across all samples in a single RNAseq dataset to prevent dominance in each principal component by high expression genes, and genes with a standard deviation of 0 across all samples within a comparison were removed. Filtering by expression prevented z-score scaled PCA from being dominated by low expression and noisy gene changes. Correlation analyses were performed between the platelet molecular maturation and functional measurements of *room-temperature stored* and *cold-stored* datasets. Further details are available in the **Supplemental Materials**.

## RESULTS

### Developing the Platelet Molecular Maturation Model

We compared two distinct endogenous platelet aging datasets (*HT*, N=8; *BB*, N=4) to determine whether RNA features at extremes of the platelet lifespan (young and old) were conserved across donors and studies. We found 6015 shared expressed genes (**Figure 1A**, Pearson’s *r*=0.761, p=2 ×10^-4^) with remarkably concordant FCs, suggesting conserved RNA metabolism across the entire platelet genome. Moreover, we identified 2008 shared differentially expressed genes (DEGs), with highly correlated FC values across datasets (**Figure 1B**, *r*=0.923, p=2 ×10^-4^). This suggests RNA metabolism is highly regulated during endogenous platelet aging. Due to the degree of concordance, moving forward, we reference the shared gene RNAseq dataset (combining *HT* and *BB)* as the *combined endogenous platelet aging* dataset, thereby defining a platelet molecular maturation spectrum in endogenous human platelets.

**Figure 1.**
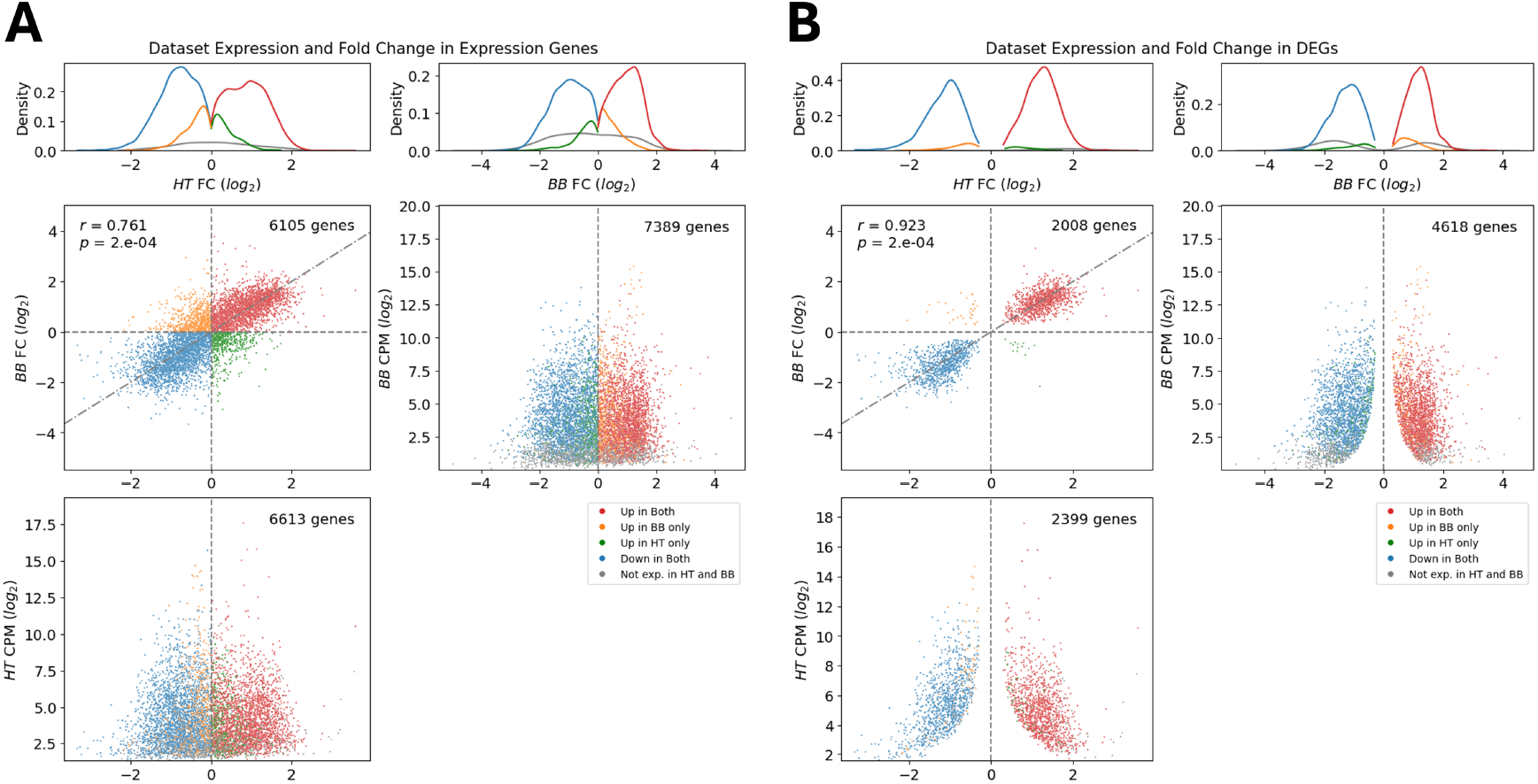
Highly regulated RNA metabolism during endogenous platelet aging provides a model of platelet molecular maturation. A. Comparison of fold change in expressed genes between FACS sorted platelet aging RNAseq datasets (“*HilleTrenk” [HT]* dataset; *“Bongiovanni-Bernlochner” [BB]* dataset). Fold change (FC) from young (high RNA content) to old (low RNA content) platelets. Coloration is determined by direction of FC (red up in both *HT* and *BB*, orange up in *BB*, green up in *HT*, blue down in both *HT* and *BB*, grey not expressed [not exp] in either) with log-2 CPM vs. log-2 FC for expressed genes in each dataset. Dashed line is a line of equivalence, not the line of best fit. Pearson correlation coefficient (r) and its significance (p) as determined by permutation are shown for each comparison. B. Comparison of FC in differentially expressed genes (DEGs) between *HT* and *BB* datasets; visualization same as in A.

### Leveraging the Platelet Molecular Maturation Model to Examine Platelet RNA Metabolism in Storage

To evaluate whether platelet molecular maturation is conserved during platelet storage, we first evaluated changes in gene expression between day 0 and 7 *room-temperature stored* datasets, then compared these findings to the *HT* and *BB* datasets, and finally to the *combined endogenous platelet aging* dataset. Gene expression FCs were highly correlated between the *HT* and *BB* datasets and the *room-temperature stored* dataset (**Figure 2A**; *HT* dataset, 5949 shared genes with *room-temperature stored* dataset, *r*=0.831, p<0.0002; **Figure 2B;** *BB* dataset, 6554 shared genes with *room-temperature stored* dataset, *r*=0.629, p<0.0002). Comparison of DEGs between *room-temperature stored* and *BB* and *HT* datasets found 1383 and 2373 shared genes, respectively (**Figure 2C**, Pearson’s *r*=0.941, p=2 ×10^-4^; **Figure 2D**, Pearson’s *r*=0.861, p=2 ×10^-4^). Gene set enrichment analysis of the *room-temperature stored* dataset compared to the *combined endogenous platelet aging* dataset showed significant enrichment of downregulated genes, but not upregulated genes (**Figure 2E; Supplemental Table 1**). This suggests that gene-specific platelet RNA metabolism, and thus, platelet molecular maturation during room-temperature storage is very similar to that seen with endogenous platelet aging.

**Figure 2.**
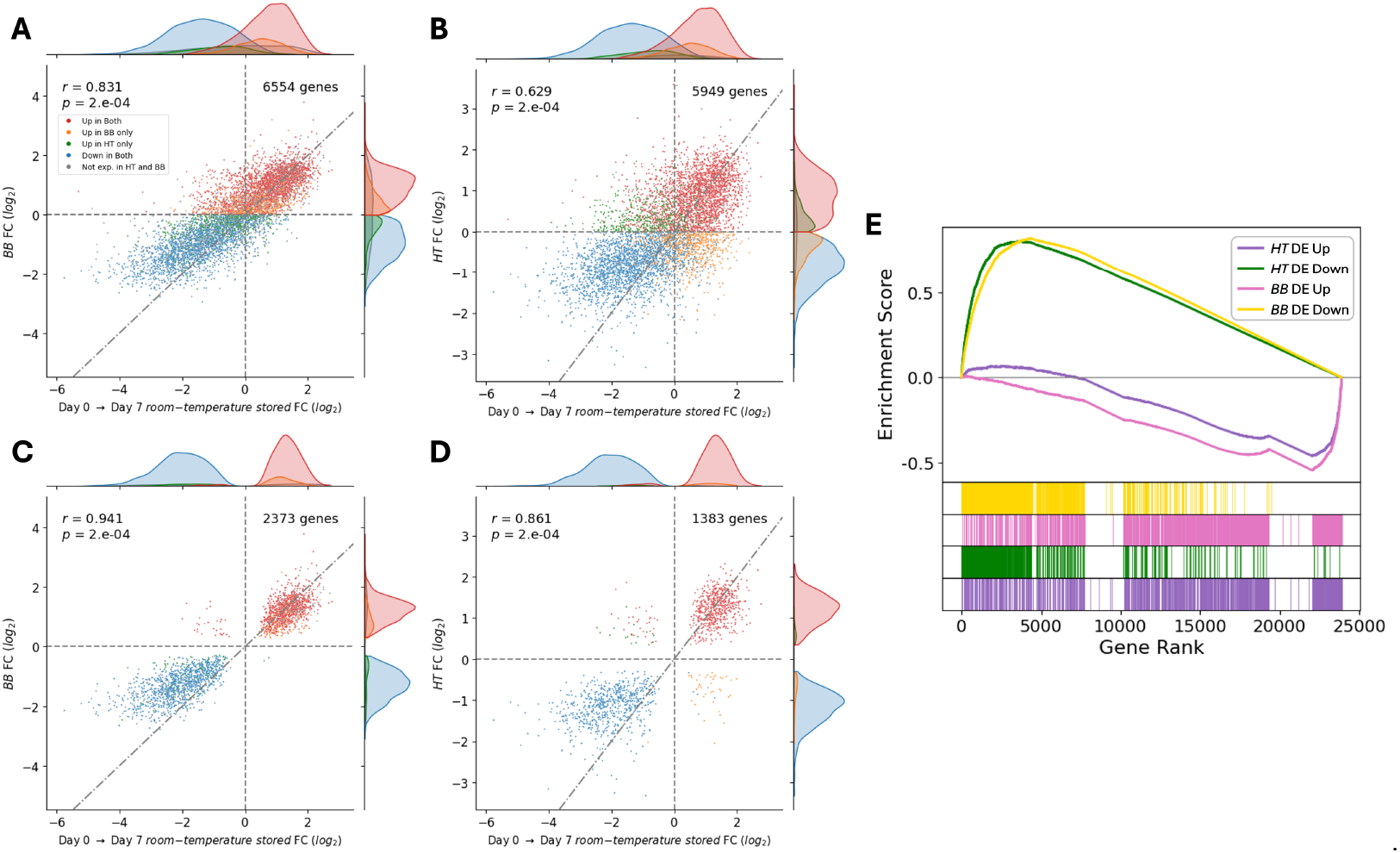
Platelet molecular maturation during room-temperature storage mirrors endogenous platelet aging. Axes represent gene expression fold change (FC) between conditions. Dashed diagonal lines show equivalent fold change (x=y), not the line of best fit. A, B. *Room-temperature stored* datasets have significantly correlated FCs in shared expressed genes with A, *BB* dataset, and B, *HT* dataset. FC is from young (high RNA content) to old (low RNA content) platelets in *HT* and *BB* datasets, and from day 0 to day 7 in *room-temperature stored* datasets. Genes were filtered for sufficient expression in datasets. To highlight the genes most similar between datasets, all genes were colored based on the direction of their FC across datasets with each quadrant of direction being colored differently (red up in both *HT* and *BB*, orange up in *BB*, green up in *HT*, blue down in both *HT* and *BB*, grey not expressed [not exp] in either). C, D. FCs with shared statistically significantly DEGs between *room-temperature stored* datasets and C, *BB* dataset, and D, *HT* dataset. Pearson correlation coefficient (r) and its significance (p) as determined by permutation are shown for each comparison E. Gene set enrichment analysis plot of *room-temperature stored* dataset expression compared to the gene sets of DEGs in the *HT* and *BB* datasets. Lower heatmaps show group membership for genes along the ranking. Above is enrichment score at each gene rank.

### Examining Age-Induced Changes to Platelet RNA Architecture

Given the enrichment in downregulated genes that we found (**Figure 2E**), we evaluated nucleotide-specific metabolism using the NLSC tool (outlined in the Methods and **Supplemental Materials**). The NLSC approach allows all significant nucleotide changes to be tallied simultaneously and compared to their position in the gene body on a 5’ to 3’ basis. We identified a preference for 3’-directed degradation in downregulated genes across *BB, HT*, and *room-temperature stored* datasets (**Figure 3A-C**). In upregulated genes, however, the number of significant NLSC position changes are minimal, with an absence of strong positional bias (**Figure 3D-F**). Collectively, these data highlight conserved 3’-directed degradation of transcripts, leading to downregulation of shared genes across the platelet molecular maturation spectrum in both endogenous and stored platelets.

**Figure 3.**
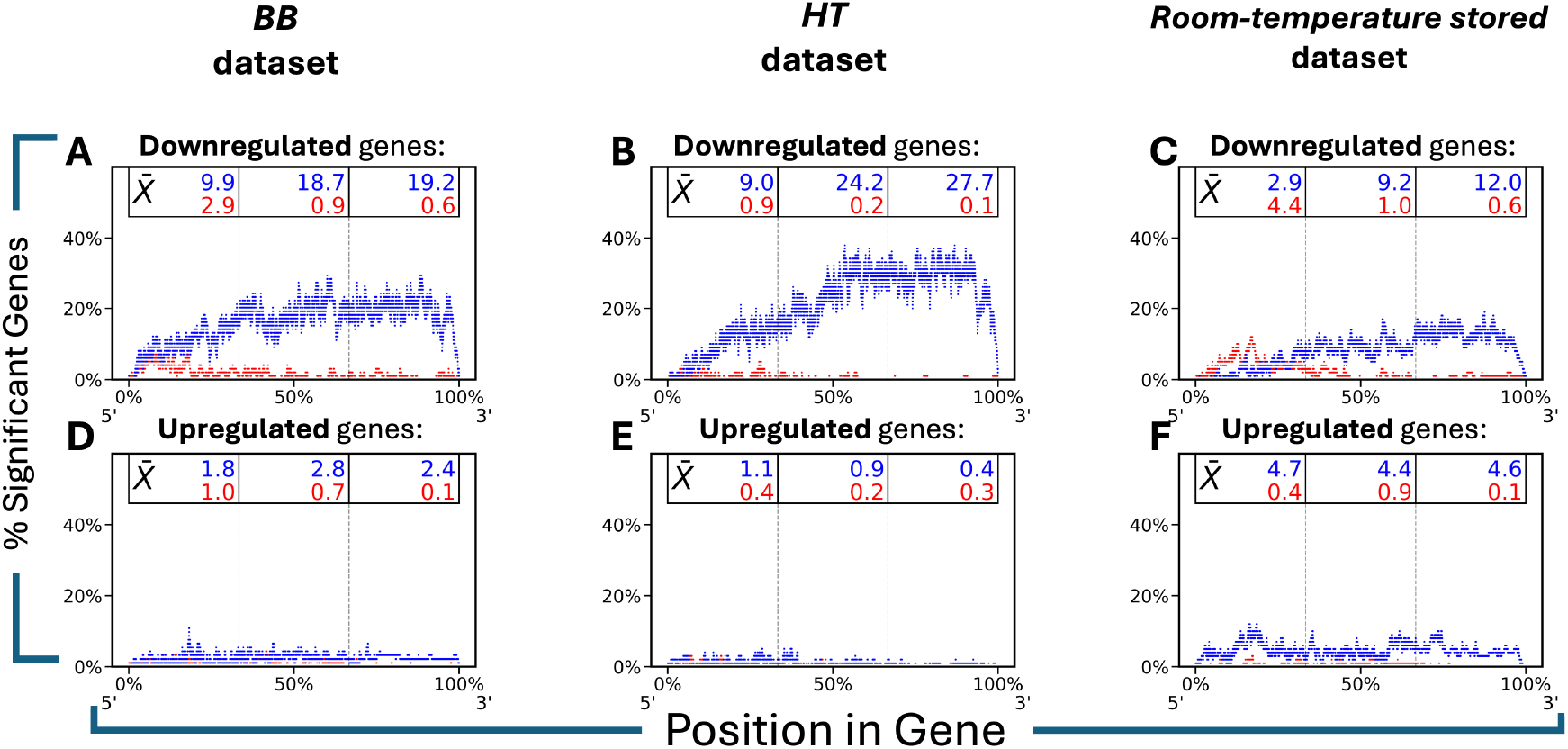
Degradation-specific RNA mechanisms across endogenous platelet aging and room-temperature storage. A-F. Condition specific Nucleotide Level Statistical Composite (NLSC) of the top 100 genes in each condition. Single genes statistically compared between groups of samples at each point. Results are tallied and displayed as a coordinate-based histogram (Blue lines represent nucleotide features going down and red lines represent features going up, relative to control). Genes were grouped by differential expression direction. A, B, C. *Negative Gene NLSC*. Limited to genes with negative expression changes. These composites reveal that 3’-directed degradation is the mechanism by which those genes are decreasing in total count. D, E, and F. *Positive Gene NLSC*. Limited to genes with positive expression changes. These composites show genes have fewer individual points that pass statistical measures and lack the structured, 3’-directed degradation.

### Identifying Conserved Genomic Patterns of Platelet Molecular Maturation

To identify conserved platelet aging RNA profiles, we performed PCA on each individual dataset (*HT, BB*, and *room-temperature stored* dataset) and the *combined endogenous platelet aging* dataset. Correlation analyses of principal components (PC) between individual datasets (**Supplemental Figure 1**) revealed strong correlation between PC1 of *HT* and *BB* (cosine distance of 0.94 with a permuted significance of *p<*0.0002; cosine distance of 0.87 with a permuted significance of *p <* 0.0002, respectively) and *room-temperature stored* dataset (cosine distance=0.62 with significance from permutation of *p*<0.0002), as well as the *combined endogenous platelet aging* dataset consistent with conserved platelet molecular maturation.

High contributing genes from the top and bottom 200 rank of the first PC loadings of the *combined endogenous platelet aging* dataset were used for gene ontology enrichment analysis (**Figure 4**). The top 200 genes are enriched for multiple cellular functions, including nucleic acid and protein metabolism (**Figure 4B**). In contrast, the bottom 200 genes represent much more specific categories of function (e.g. aggregation, cytoskeleton, and vesicular trafficking) with significantly higher fold enrichment and better false discovery rate corrected p-values (**Figure 4C**). As noted in **Figure 3**, downregulated genes are specifically targeted for 3’-directed degradation while upregulated genes increase ‘passively’ increasing secondary to the read displacement phenomena, described in **Supplemental Materials** (**Supplemental Figure 2D**). Taken together, these data are strongly suggestive of a regulated RNA metabolism that selectively degrades genes involved in platelet function and hemostatic activity during endogenous platelet aging and storage.

**Figure 4.**
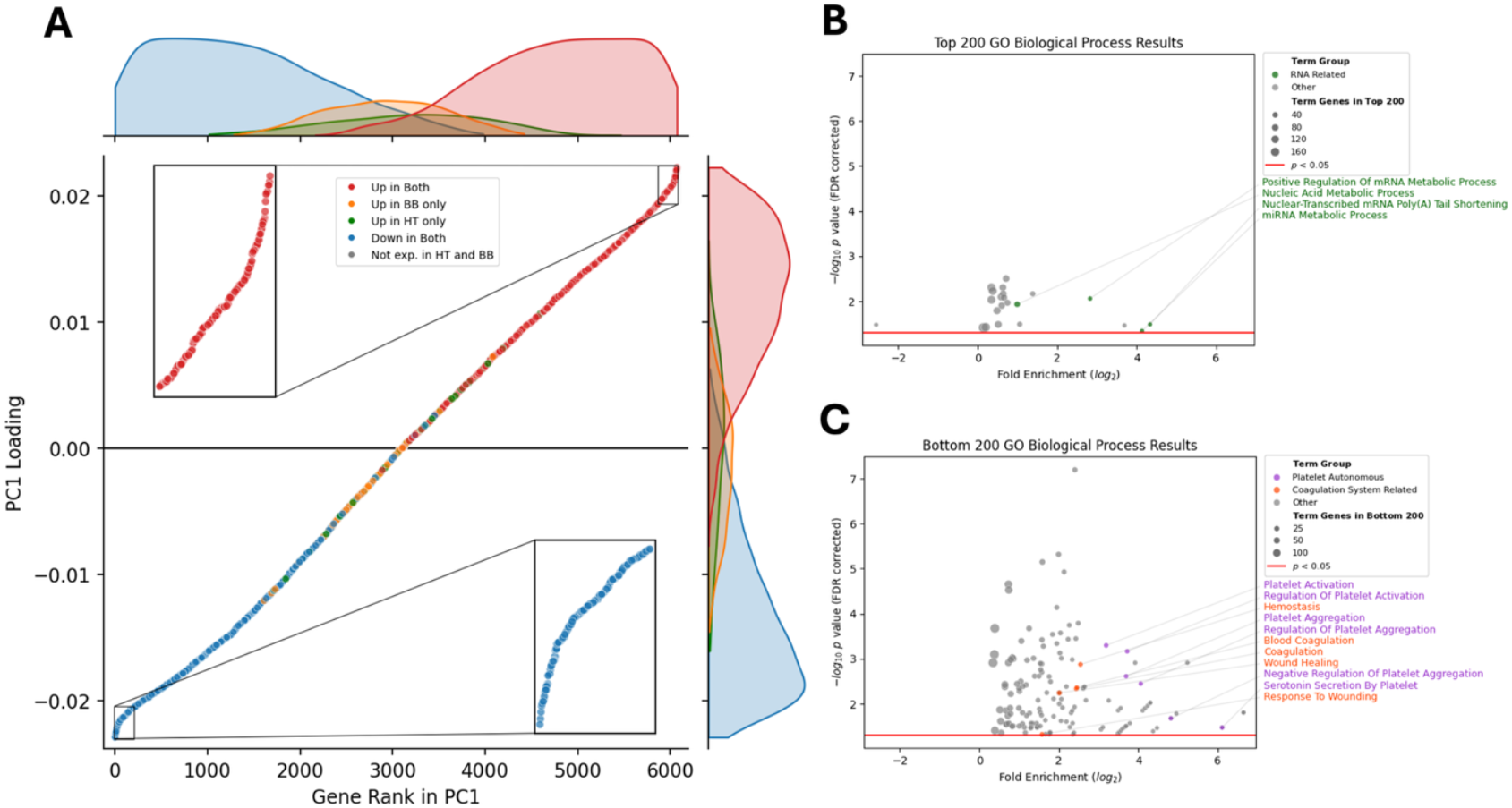
Conserved platelet molecular maturation across endogenous platelet aging and room-temperature storage. A. The first principal component (PC1) of *combined endogenous platelet aging* dataset plotted by gene loading against gene loading rank. Color is derived from fold change direction in each dataset as per Figure 1-3. Insets highlight the top and bottom 200 genes used in gene ontology enrichment analysis. B, C. PANTHER Gene Ontology statistical overrepresentation results for the 200 most positively contributing genes (B) and 200 most negatively contributing genes (C) to PC1 of the *combined endogenous platelet aging* dataset. Horizontal axis denotes the log fold enrichment of genes found with GO term relative to the expected number. Vertical axis denotes the negative log-10 FDR correct p value. Size of circle relates the number of genes found in PC1 matching that GO term. Color here denotes classes of identified GO terms which were determined by keywords. See **Supplemental Sheet 1** for full results.

### Defining Relationships of Platelet Molecular Maturation and Platelet Function

To understand platelet function in the context of platelet molecular maturation, we first measured hemostatic function across storage. As previously identified^25-28^, we confirmed cold-storage preserved platelet aggregation over time (**Figures 5A,B**), as well as retained thrombin generating capacity (**Figure 5C**) and maximal clot formation in viscoelastic testing (**Figure 5D**) when compared to room-temperature stored platelets. Next, we evaluated which platelet functional metrics correlated the most strongly across storage temperatures and duration as a stable signature of platelet functional variation (**Figure 5E**). We then correlated these platelet functional measurements with genomic signatures. Maximum amplitude in aggregation response induced by ADP and epinephrine [ae] or ADP and collagen [ac] was projected against our platelet molecular maturation scale, which yielded strong correlation values (**Figure 5F**; *r*= - 0.88 and -0.66, respectively). Thus, the platelet molecular maturation range is correlated with aggregation capacity.

**Figure 5.**
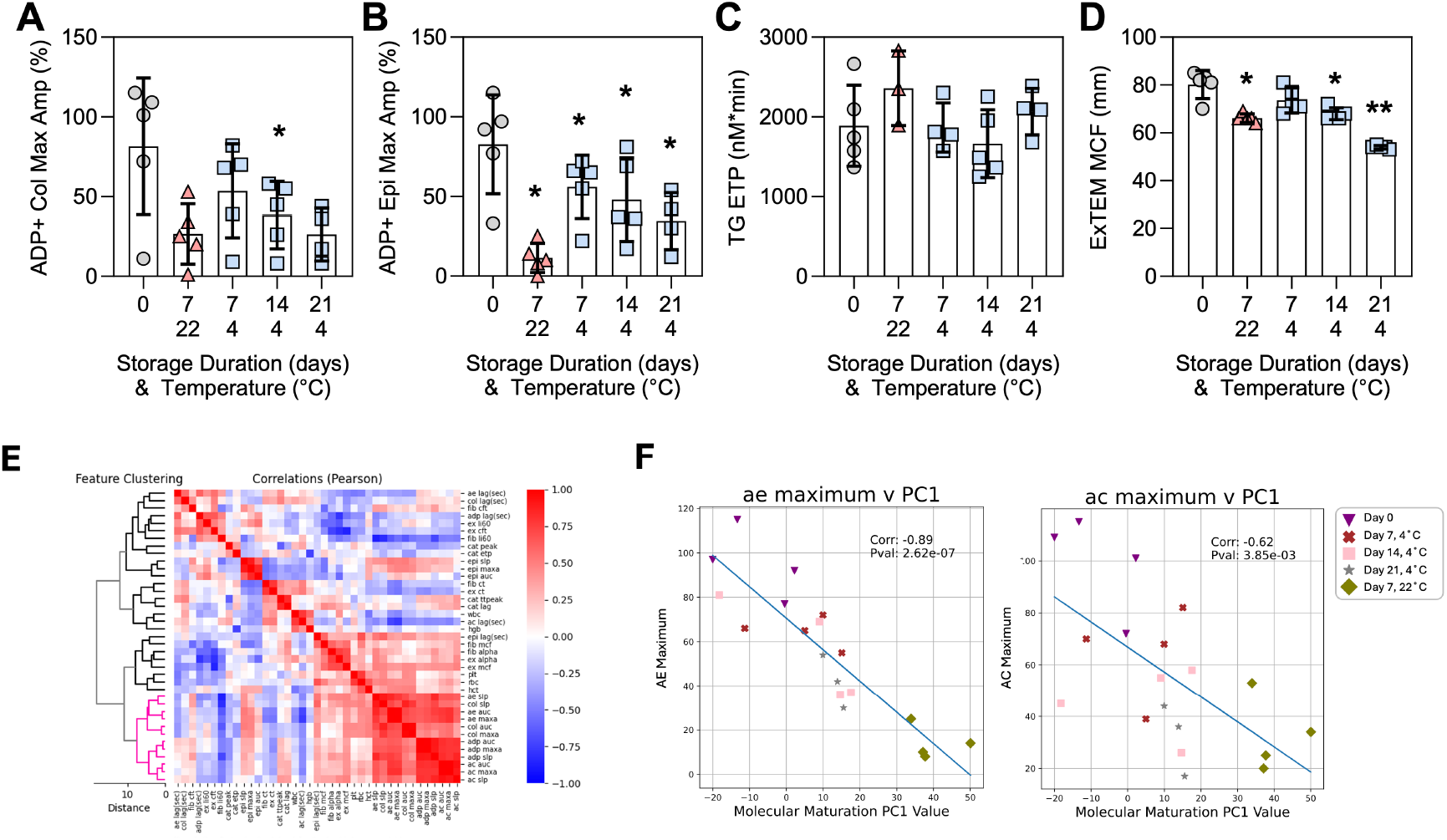
Correlation of platelet molecular maturation and platelet function. A-D. Stored platelet functional measurements (N=3-5 for each temperature and timepoint); one way ANOVA, with each timepoint compared to baseline (Day 0); *, p<0.05; **, p<0.01. A. Maximum amplitude of aggregation of stored platelets stimulated by adenosine diphosphate (ADP) and collagen (Col), and B. ADP an epinephrine (Epi). C. Endogenous thrombin potential (TG ETP), and D. Maximum clot firmness in response to rotational thromboelastometry extrinsic activation (EXTEM MCF) E. Recursive Euclidean clustering of all z-score normalized tested functional measurements produced a core set of features that are most consistently altered during platelet storage (pink cluster). Heatmap is colored by the Pearson’s correlation coefficient between each functional measure (see **Supplemental Table 2**). F. Functional measurements compared against projection of *room-temperature stored* and *cold-stored* datasets expression into principal component 1 (PC1) of the *combined endogenous platelet aging* dataset for all functional measurements (see **Supplemental Figure 5**). Examples of two scatter plots showing strong correlation between projection into the *combined endogenous platelet* aging PC1 and platelet aggregation function are shown. Measurement performed by light transmission aggregometry ADP and epinephrine (ae) maximum amplitude and ADP and collagen (ac) maximum amplitude (correlation coefficients of -0.88 and -0.66, respectively). Time and temperature condition color scheme is noted to the right.

### Uncovering Effects of Storage Temperature on Platelet RNA Metabolism

Because cold-storage preserves platelet hemostatic activity, and our platelet molecular maturation model identifies genomic profiles associated with stored platelet function, we hypothesized that cold-storage-induced alterations in RNA metabolism underlie the preserved hemostatic function of cold-stored platelets. Indeed, we found that cold-storage substantially slows the kinetics of platelet RNA metabolism over time (**Figure 6**). Specifically, there is significant divergence between day 0 and day 7 *room-temperature stored* datasets. In contrast, no significant differences were observed between day 0 and days 7, 14, or 21 of *cold-stored* datasets. This effect is reflected in the compressed projection range of *cold-stored* datasets relative to *room-temperature stored* datasets (**Figure 6A, B; 3^rd^ panel**). Moreover, using our NLSC analysis, we found that cold-storage significantly attenuates 3′-directed degradation compared with room-temperature storage (**Figure 6C**). In contrast to the pronounced asymmetric, 3′-directed degradation observed during room-temperature storage (**Figure 3**), RNA degradation under cold-storage is markedly less asymmetric and occurs at substantially lower levels (**Figure 6C**). The proportion of significant genes within the terminal third of transcripts decreases from 19.4% in platelets at room-temperature storage for 7 days to 0.5%, 1.6%, and 3.5% after 7, 14, and 21 days of cold-storage, respectively. The modest increase observed over prolonged cold-storage likely reflects non-specific degradation occurring elsewhere within coding sequences (data not shown), consistent with the inherent instability of RNA in anucleate cells.

**Figure 6.**
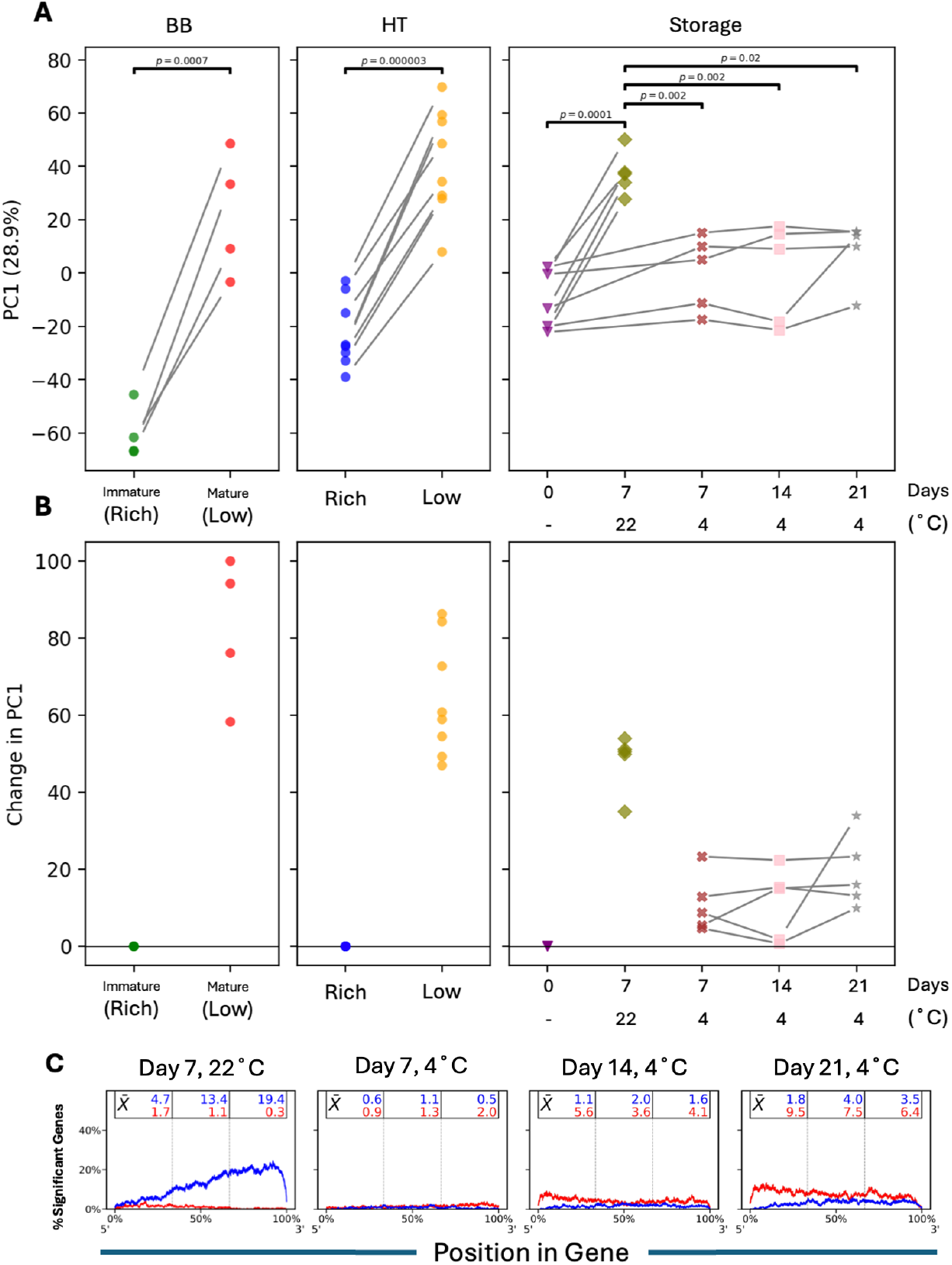
Cold-storage slows platelet RNA metabolism and decreases 3’-directed degradation. A. *HT, BB*, room-*temperature stored*, and *cold-stored* projection into *combined endogenous platelet aging* dataset principal component 1 (PC1). Grey lines indicate pairing between platelet samples isolated together. Statistical comparisons by ANOVA with a Tukey’s Honestly Significant Difference post-hoc correction for *room-temperature stored* dataset and *cold-stored* dataset comparisons; T-test for *HT* and *BB* dataset comparisons. Significant differences denoted by p values. B. Projection into *endogenous platelet aging* dataset PC1 with samples normalized to control (young for *HT* and *BB* datasets, and to day 0 for *room-temperature stored* and *cold-stored* datasets). C. Nucleotide based counting demonstrates cold-storage significantly attenuates 3′-directed degradation compared with room-temperature storage. Condition specific Nucleotide Level Statistical Composite (NLSC) of the top 500 genes by expression. Blue lines represent nucleotide features going down and red lines represent features going up relative to control. All samples are compared to baseline (Day 0) and labeled by room-temperature storage (22°C) at day 7, and cold-storage (4°C) at days 7, 14 and 21 days, respectively.

Analysis of differential expression revealed a conserved set of genes throughout cold-storage as compared to room-temperature storage (**Figure 7 and Supplemental Figure 4**). A total of 669 genes were differentially expressed between *room-temperature stored* and *cold-stored* datasets (**Figure 7A-C**). In contrast, no differentially expressed autosomal genes were detected with any length of cold-storage (**Figure 7D inset**), indicating persistent transcript stability during cold-storage. Gene ontology analyses revealed that transcripts dysregulated during room-temperature storage are enriched for aging-associated pathways, whereas cold-storage maintains a transcript profile more closely resembling that of young platelets (**Figure 7E, F**). Notably, inter-individual genotypic variation appears to contribute more prominently to platelet preservation in cold-storage than during room-temperature storage (**Figure 6A, B; 3^rd^ panel; Supplemental Figure 5**).

**Figure 7.**
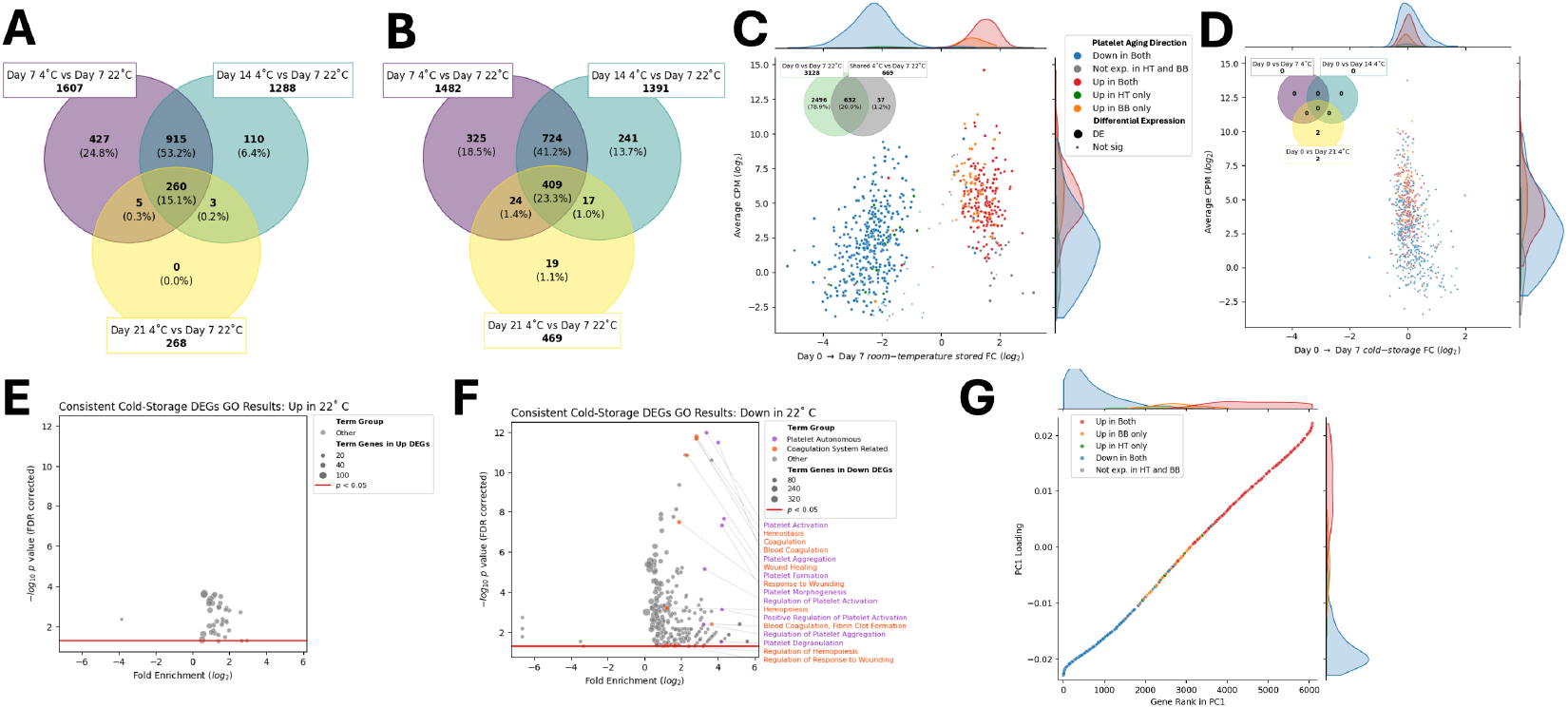
Cold-storage preserves platelet RNA relative to room-temperature storage. A. Overlap of genes with higher expression in day 7 *room-temperature stored* dataset compared with *cold-stored* datasets across timepoints; 260 genes are shared across all comparisons. B. Overlap of genes with lower expression in day 7 *room-temperature stored* dataset compared with *cold-stored* datasets across all timepoints; 409 genes are shared across all comparisons. C. Fold change (FC) versus expression of the 669 genes consistently differentially expressed between day 7 *room-temperature stored* dataset and *cold-stored* datasets across all timepoints (intersection of A and B), shown for day 7 *room-temperature stored* dataset. The inset compares these 669 genes with all genes differentially expressed between day 0 and day 7 *room-temperature stored* datasets. Gene colors reflect FC direction in *HT* and *BB* datasets, and symbol color and shape denote differential expression status. D. FC versus expression of the same 669 genes for day 7 *cold-stored* dataset. The inset compares genes differentially expressed between day 0 and all *cold-stored* datasets. Gene colors and symbols are defined as in (C). The two differentially expressed genes are mitochondrial genes (*MT-CO2* and *MT-CO3)* in the day 21 *cold-stored* dataset. E. Gene ontology overrepresentation analysis of the 260 genes with higher expression in day 7 *room-temperature stored* relative to *cold-stored* datasets. F. Gene ontology overrepresentation analysis of the 409 genes with lower expression in day 7 *room-temperature stored* relative to *cold-stored* datasets. G. Expression of 637 of the 669 consistent genes detected in *HT* and *BB* datasets plotted against PC1 of the *combined endogenous platelet aging* dataset.

## DISCUSSION

Changes in transcriptomic expression are synonymous with cell fate, yet platelet biology has not found an indisputable role for intrinsic RNA expression.^16,42-49^ In this study, we sought to define the central molecular features that govern platelet aging, and to determine whether these processes also drive the functional changes characteristic of platelet storage. By integrating multiple high-quality RNAseq datasets with novel analytic approaches tailored to the biology of anucleate platelets, we show that platelet aging is not a passive process of transcript decay, but rather follows a structured and reproducible molecular trajectory both endogenously and in storage, which we term platelet molecular maturation. Storage temperature emerged as a dominant modifier of this trajectory, with cold-storage markedly slowing RNA metabolic kinetics and preserving transcripts associated with younger, more hemostatically competent platelets. Together, these findings provide mechanistic insight into the platelet storage lesion and identify transcriptomic features that may serve as biomarkers or therapeutic targets to extend platelet shelf life.

Our first major finding is that platelet aging follows a conserved program of regulated RNA metabolism. Although platelets are anucleate and incapable of de novo RNA synthesis, they retain a complex RNA repertoire capable of stimulus-dependent translation, supporting the concept that post-transcriptional mechanisms shape platelet function and that platelet RNA profiles change with activation and aging, implicating selective RNA processing and degradation as contributors to platelet heterogeneity and functional adaptation.^15,16,48,50^ Across two independent datasets of young and old human platelets, we observed striking concordance in gene expression, differential expression patterns, and RNA structural features, demonstrating that the platelet transcriptome evolves in a coordinated and non-random manner throughout the lifespan. Nucleotide-level analyses revealed selective and directional RNA processing, with preferential 3′-directed degradation of downregulated transcripts. This pattern was conserved across endogenous aging and room-temperature storage, arguing against stochastic RNA decay as the primary driver of transcriptome remodeling. Instead, these data support a model of enzymatically regulated RNA metabolism, potentially mediated by exonucleases and RNA-binding proteins. The enrichment of downregulated transcripts involved in platelet-autonomous functions, including aggregation responses, cytoskeletal dynamics, and vesicular trafficking, suggests that platelet RNA metabolism actively tunes platelet functional capacity over time. Transcripts appearing upregulated were largely explained by displacement phenomena arising from selective degradation of other transcripts rather than increased expression. These observations refine our understanding of the post-transcriptional regulation of platelets by demonstrating that platelet aging is a regulated process involving targeted RNA decay that shapes the platelet’s evolving phenotype.

Second, we demonstrated that platelets in room-temperature storage recapitulate the transcriptomic trajectory observed during endogenous platelet aging. Gene expression changes in stored platelets aligned closely with datasets of young and old platelets, and principal component analyses confirmed that storage-associated shifts in platelet RNA metabolism fall along the same molecular continuum as endogenous platelet aging. These results provide a mechanistic explanation for similarities between old circulating platelets and room-temperature stored platelets, including diminished aggregation responses, reduced granule secretion, and altered metabolic capacity.^14^ The accelerated progression along the platelet molecular maturation axis during room-temperature storage suggests that the platelet storage lesion represents an exaggeration of the natural aging process, governed by RNA metabolism.

A third major insight is that platelet molecular maturation strongly correlates with platelet functional outputs. Aggregation responses, thrombin generation, and viscoelastic clot formation measurements tracked closely with a sample’s position along the platelet molecular maturation spectrum. These findings provide direct evidence that transcriptomic aging is functionally meaningful, even in anucleate cells, and are consistent with findings by Hille et al.^15^ showing greater responsiveness of young platelets compared to non-reticulated platelets. These findings extend prior observations that young and old platelets differ in reactivity and metabolic capacity^14^, and suggest that the platelet transcriptome is a dynamic regulator of platelet behavior throughout the platelet lifespan. Platelets harbor a complex RNA landscape, including pre-mRNAs, mature mRNAs, and regulatory non-coding RNAs, which can be selectively processed and translated in response to activation and extracellular stimuli, allowing platelets to rapidly modify their proteome in health and disease.^43^ This dynamic RNA regulation contributes to platelet heterogeneity and functional plasticity in hemostasis, inflammation, and immune responses, and our data further support that regulatory dynamics are also central during platelet storage.

Perhaps the most translationally significant and novel insight from this study is that cold-storage profoundly alters the kinetics of platelet molecular maturation. Cold-stored platelets exhibited a compressed molecular aging trajectory, attenuated 3′-directed degradation, and remarkable transcript stability across extended storage durations. No autosomal genes were differentially expressed with increasing cold-storage duration, indicating near-complete arrest of transcriptomic aging under these conditions. In contrast, hundreds of genes diverged between cold and room-temperature storage, with room-temperature stored platelets selectively enriching for aging-associated pathways. These findings provide a mechanistic basis for the preserved hemostatic function and relative hyperreactivity observed in cold-stored platelets. Reduced activity of RNA exo- and endonucleases at cold temperatures offers a plausible mechanism for this preservation, although direct interrogation of nuclease activity will be required.

Cold-stored platelets also maintained transcriptomic profiles more similar to early stages of platelet molecular maturation, suggesting that cold storage effectively pauses platelet aging rather than inducing an alternative stress response. This reconciles longstanding observations of superior clot formation and thrombin generation in cold-stored platelets while explaining their altered behavior in inflammatory or immune contexts. Notably, cold-storage appeared to unmask inter-individual genotypic variability in platelet preservation, whereas room-temperature storage imposed a dominant and uniform aging program. This observation aligns with emerging data implicating donor-specific effects in cold-stored platelet behavior.^51^ These findings raise the possibility that molecular profiling could identify donors whose platelets are particularly well suited for extended cold-storage or specific clinical indications, enabling more personalized transfusion strategies.^52^

Collectively, our findings reshape the understanding of platelet aging and storage. Rather than viewing stored platelets as a system undergoing nonspecific and non-physiologic degradation, our results position them within a broader continuum of physiologic aging governed by conserved platelet RNA metabolic processes. The platelet storage lesion, long defined phenotypically^53-57^, can now be conceptualized at the transcriptomic level, enabling identification of biomarkers that more accurately reflect platelet quality than storage duration or current limited measures alone. These biomarkers could improve inventory management, reduce wastage, and identify units with higher functional potential. Understanding regulated RNA metabolism in platelets also opens the possibility of targeted interventions to preserve transcripts associated with hemostatic function. For example, inhibiting specific 3′-to-5′ exonucleases or modulating RNA-stabilizing proteins might slow platelet molecular maturation and prolong the associated platelet function of room-temperature stored platelets. As cold-storage re-emerges as a viable strategy, our data provide mechanistic support for its hemostatic advantages and clarify the genomic basis for its divergence from room-temperature storage.

Several limitations warrant acknowledgment. While our analyses implicate RNA metabolism as a driver of platelet aging and storage-associated changes, the specific enzymes involved remain undefined. Additionally, although we demonstrate strong correlations between platelet molecular maturation and functional behavior, causal relationships between individual transcripts and platelet function will require targeted perturbation studies. Finally, the clinical implications of prolonged cold-storage, including effects on platelet clearance, immune interactions, and post-transfusion survival, were not addressed.

This study raises important questions for future investigation, including defining the enzymatic machinery driving platelet RNA metabolism, exploring links between platelet RNA metabolism and non-hemostatic platelet functions, and extending the platelet molecular maturation model to disease states. Prospective application of this model in transfusion medicine may clarify whether transcriptomic markers predict post-transfusion recovery or efficacy. Given the growing platelet supply crisis in the US, the development of high-fidelity molecular biomarkers for platelet quality could have enormous clinical impact.^24,58-61^

In summary, this work establishes platelet molecular maturation as a conserved and functionally relevant model for understanding platelet aging in physiologic and storage contexts. By demonstrating that cold-storage dramatically slows platelet RNA metabolic aging and preserves a youthful platelet transcriptome, our findings provide a unifying molecular explanation for the preserved hemostatic function of cold-stored platelets. More broadly, this study introduces a platelet genomic atlas that can inform biomarker development, optimize storage strategies, and guide the design of next-generation platelet products with improved efficacy and reduced waste.

## Supporting information

Supplemental Sheet 1

## ACKNOWLEDGEMENTS

Funding: Research reported in this publication was supported by the National Institute of General Medical Sciences of the National Institutes of Health under Award Number R35GM150656 (LZK), and by an award from the American College of Surgeons (LZK). SMS is supported by 5K25HL161401. KAT is supported by R01HL171594. The content is solely the responsibility of the authors and does not necessarily represent the official views of the National Institutes of Health or of the American College of Surgeons.

## AUTHORSHIP CONTRIBUTIONS

L.Z.K. and R.J.B. and conceived and designed the study. L.Z.K., R.J.B., A.T.F., K.Z., N.M., F.M., S.M.S., J.M.M., and K.C.R., performed experiments and collected data. R.J.B., C.M.V.B., Y.A.S., S.M.S., J.M.M., and K.C.R. conducted bioinformatic and statistical analyses. F.M., R.J.B., J.S., D.W., G.C., C.M.V.B., Y.A.S., A.T.F., K.Z., L.Z.K., N.M., S.M.S., J.M.M., K.C.R., and K.T. interpreted the data. L.Z.K., R.J.B., C.M.V.B., S.M.S., and K.T. drafted the manuscript. All authors critically revised the manuscript for important intellectual content, approved the final version, and are accountable for all aspects of the work.

