## Supplemental Materials for "Platelet Molecular Maturation Links Platelet Aging and the Platelet Storage Lesion"

### SUPPLEMENTAL METHODS

Additional Details of the Collection, Functional Measurements, and Sequencing of Stored Platelets

Platelet concentrates (N=5) were collected from healthy volunteer donors (University of Pittsburgh Institutional Review Board #21110093) via apheresis using a Trima Accel 7 (Terumo, Somerset, NJ). Platelets were resuspended in autologous plasma and ACD-A in accordance with U.S. Food and Drug Administration regulations and American Association for Blood and Biotherapies standards and guidelines. Units were then split into equal volumes using the attached transfer bag, and one unit was stored at 22°C (room-temperature stored; agitated), and the other at 4°C (cold-stored; unagitated) per current standard practice. A baseline sample collected from the starting bag prior to splitting was isolated (day 0), and units were aseptically sampled weekly: room-temperature stored units were isolated on day 7 (current maximum expiry) and cold-stored units isolated on day 7, day 14, and day 21 (maximum potential shelf life). Complete blood counts were performed using an Advia 2120i (Siemens, Tarrytown, NY), residual white blood cells were measured using the ADAM rWBC2 (NanoEntek, Seoul, Korea), and hemostatic functional testing was performed as described below. While apheresis platelets are leukoreduced by the collection process, they can still contain residual leukocytes as well as residual red blood cells. We therefore subjected the stored platelet samples to purification procedures described below prior to sequencing.

Platelet hemostatic assessment was performed using a broad panel. Light transmission aggregometry was performed using a Model 700 Whole Blood/Optical Lumi-Aggregometer (ChronoLog Corporation) with single agonism (adenosine diphosphate [ADP], epinephrine, collagen) and dual agonism (ADP+epinephrine, ADP+collagen). Lag time, slope, maximum amplitude, and area under the curve were recorded. Thrombin generation was measured using a Calibrated Automated Thrombogram (CAT; Stago, Parsippany, NJ) with the platelet-rich plasma (PRP) reagent. Lag time, peak thrombin, and endogenous thrombin potential (ETP) were recorded. Viscoelastic testing was performed using a rotational thromboelastometry (ROTEM) delta (Werfen, Bedford, MA) with EXTEM (extrinsic cascade) and FIBTEM (extrinsic cascade with platelet inhibition) reagents per device and manufacturer instructions. Clotting time, clot formation time, maximum clot formation, and lysis index at 60 min (LI60) were recorded.

Samples from stored platelets were incubated with PSG and PG-E1, and spun at 800x*g* (20 min, room temperature, no brake), resulting in a platelet pellet. Supernatant was again removed and additional PSG and PG-E1 was added to the pellet, which was then gently resuspended. The sample was then mixed with anti-CD45 beads and sorted by magnetic column. After column purification, the sample was centrifuged again at 800x*g* in the same fashion, and the resulting pellet resuspended in Trizol and stored at -80°C. From platelet lysates stored in Trizol, RNA was isolated using the Qiamp RNA Blood Mini Kit (Qiagen, Hilden, Germany). Deoxyribonucleic acid (DNA) contamination was removed with DNAse I, and sample quality was assessed by Bioanalyzer (Agilent, Santa Clara, CA). Complementary DNA libraries were generated with 1 ng to 2 ng of RNA using the Ovation random primed isothermal amplification system (NuGen, Redwood City, CA). Sequencing was performed at a depth of 100 to 400 million reads using a NovaSeq 6000 (Illumina, San Diego, CA).

Additional Details of the Sequence Alignment and Mapping

For all RNA sequencing (RNAseq) data (public repositories and stored platelets), quality was assessed with several tools. First, general statistics were measured and compared with FastQC[34]. Next, read duplication was counted with the RSeQC package[35]. We used the Picard software suite from the Broad Institute[36] to measure mapping classifications (coding, UTR, intronic, intergenic or ribosomal) for all mapped reads. Furthermore, all quality control metrics were visualized with MultiQC[37]. RNAseq data from all sources was aligned to the GRCh38 assembly of the human genome and mapped with release 100 of the Ensembl gene annotations using the STAR aligner (version 2.6.0c)[38]. Gene expression values were counted using featureCounts from the Subread package[39], considering genes annotated in the previously mentioned GRCh38 assembly.

Additional Details of the Comparative Differential Gene Expression Analysis

For each RNAseq dataset, gene expression for control groups were compared to experimental with edgeR[40]. Gene counts were normalized by trimmed mean of M values and genes were filtered for expression by *filterByExpr* as implemented by edgeR[62]. Gene expression was fit with a Negative Binomial Linear Model and fit by a Quasi Likelihood test. Expression statistics for all datasets was performed in a paired manner for common donors between experimental condition. False discovery rate correction was performed for significance using the Benjamini-Hochberg correction and a false discovery rate of less than 0.05.

Genes passing expression minimums in datasets were plotted by log base-2 fold change from the high RNA condition to the low RNA condition or from the baseline (day 0) to condition. Correlation of fold change was examined by Pearson correlation coefficient and significance of correlation was assessed by permutation. Permutation was chosen for significance testing because it is robust to non-Gaussian and heteroskedastic distributions which were introduced by filtering for differential expression. Plotting and calculations were performed with the Python language, taking advantage of the *pandas*, *matplotlib*, and *seaborn* libraries[63-65]. Permuted significance was estimated with the *scipy* library *permutation_test* function using *pairings* with default parameters.[66] A platelet molecular maturation model was developed using existing split-sample RNAseq datasets of fluorescence-activated cell sorting (FACS) sorted platelets at the ends of the platelet lifespan ("Hille-Trenk"[15] *[HT]* dataset and "Bongiovanni-Bernlochner"[16] *[BB]* dataset).

Genes were colored across all figures based on the direction of the gene fold change between *HT* and *BB* datasets. Coloration by direction of fold change across both RNAseq datasets was selected as a simple visualization tool of which genes have a consensus direction in FACS sorted endogenous platelet aging experiments (red and blue) and which genes are less consistent (yellow and green).

Gene set enrichment analysis was performed on the CPM normalized gene expression of the day 0 and day 7 *room-temperature stored* datasets after the removal of mitochondrial genes. Differentially expressed genes from HT and *BB* datasets were separated by fold change direction and used as gene sets. Gene set enrichment analysis was performed with the *GSEA* module of the *GSEApy* Python library[41].

Additional Details of the RNA Metabolism Analysis

The anucleate nature and long half-life of platelets required new tools to understand RNA metabolism during platelet aging. In brief, this required the construction of a gene coverage map down to the nucleotide level and multiple data aggregation steps to elaborate the nature of intragenic RNA processing events, outlined in **Supplemental Figure 2A; Supplemental Figure 3A.** For the purposes of this paper, we focus on RNA metabolism as a summation of the statistically differing individual nucleotides gained and lost over maturation to examine what regions had the most coverage changing events. We have termed this a "Nucleotide Level Statistical Composite" (NLSC).

A nucleotide level statistical composite counts bulk nucleotide shifts that can be segregated by different criteria. For example, differential expression genes are separated by up and down regulation, and those gene classes can be further partitioned into negative or positive changed positions. This is a particularly useful visual for considering enzyme 5' or 3' location of nucleotide processing and provides clues to the different types of RNA metabolism. A cumulative score for an NLSC range is broken into three bins, a 5', a middle, and 3' bin. A relative polarization ratio can be calculated by dividing the 3' most bin by the 5' most bin to obtain the targeted coverage losses.

Additional Details of Integrated Data Analyses

In order to capture the relationships between gene expression changes, principal component analysis (PCA) was performed. Gene expression was normalized to CPM after the removal of mitochondrially derived transcripts. Genes included for this analysis follow the same expression criteria as our differential expression analysis. Specifically, genes whose expression meet the criteria described by Chen et al.[67] as implemented by *edgeR*. PCA was applied to the datasets.

Gene expression was z-score normalized across all samples in a single RNAseq dataset to prevent dominance in each principal component by high expression genes. In the RNAseq dataset combining both platelet aging experiments (*HT* and *BB* combined as the *combined endogenous platelet aging* dataset), z-score normalization was performed after RNAseq dataset combination. Genes with a standard deviation of 0 across all samples within a comparison were removed. Filtering by expression (see *filterByExpr* above for exact implementation) prevented z-score scaled PCA from being dominated by low expression and noisy gene changes potentially due to expression changes at the lower limit of detection. Normalization and filtering were performed with the Python library *pandas* and PCA was performed with the *scikit-learn* library.

Principal components were compared between RNAseq datasets by examining the absolute cosine distance between the top components. Genes which had been removed from one PCA due to a standard deviation of 0 were removed prior to the cosine distance calculation. Calculations were performed with the *pandas* library and plotting were performed with the *seaborn* and *matplotlib* libraries in Python. The first principal component appeared to be the most consistent between the three different PCAs on the *HT* and *BB* datasets. The second principal component of the *combined endogenous platelet aging* dataset appeared to be dominated by a batch effect, while components three and beyond seemed dominated by similarity to the *HT* dataset and included less of the *BB* variation. The first principal component of the *combined endogenous platelet aging* dataset was selected as an estimate of the greatest variability that occurs consistently in platelet aging experiments. This component also demonstrated the greatest similarity of any principal component in platelet aging to variability in platelet storage conditions, though notably lower similarity to than to the other platelet aging datasets.

The significance of cosine distances was determined through permutation analysis. Gene loadings were shuffled such that loading values were paired to a random gene across principal components, and then cosine distances were calculated. Significance was determined by the percentile of the observed cosine distance relative to shuffled distances. For each comparison 9,999 permutations were calculated, and significance was estimated in a two-tailed manner. Calculations relied on the Python *pandas* library and permutations were generated with the *scipy permutation_test* function.

The first principal component of the *combined endogenous platelet aging* dataset was examined and colored by platelet aging quadrant. Genes were ranked by contribution to the first principal component (also referred to as "loading") and the top and bottom 200 genes were examined by PANTHER DB for statistical overrepresentation in a gene ontology biological process. Platelet aggregation was found to be high in the positive component. The negative component demonstrated no platelet signature, suggesting a potential for reads shifting up in expression due to the loss of platelet integrin features.

*Room-temperature stored* and *cold-stored* datasets were projected into the principal component space developed from the *combined endogenous platelet aging* dataset. Stored sample expression was CPM-normalized after the removal of mitochondrially derived transcripts. Expression was filtered only to genes that passed filtering for the PCA of the *combined endogenous platelet aging* dataset. To match the z-score normalization from model training, expression across the *combined endogenous platelet aging* dataset was transformed by subtracting the mean and dividing by the standard deviation from the expression of the *room-temperature stored* and *cold-stored* datasets. Finally, sample projection was performed by multiplying the transformed expression by the loadings of principal components from PCA trained on the *endogenous platelet aging* dataset. Projected stored platelet samples were compared to the first principal component of the platelet aging dataset. All conditions were found to be statistically significantly different between day 0 and day 7 *room-temperature stored* datasets by a pairwise Tukey Honestly Significant Difference test ($p < 0.05$). No other conditions had differing projected values. Calculations were performed with the Python library *statsmodels*. Plotting and data manipulation relied on the *pandas*, *seaborn*, and *matplotlib* libraries.

The functional measurements of the stored platelets were clustered with the *scikit-learn* package[68] by performing recursive Euclidean clustering to minimize the variance of clusters being merged. To visualize this clustering, a heatmap showing individual correlations between each functional metric was produced with *seaborn*[63] and *scipy*[66]. The strongest resulting cluster was used as our core feature set to which different versions of platelet molecular maturation were compared. All functional metrics that were a member of the strongest cluster were then correlated against the platelet molecular maturation score for each of the platelet storage conditions, separately by individual donor. The aggregated results of those correlation coefficients were used to score each training set for the platelet molecular maturation scale. Finally, functional assessments were correlated against the selected platelet molecular maturation scale and visualized, performed with *scipy* and *matplotlib*[65,69].

**Supplemental Table 1. Gene set enrichment analysis results.** Gene set enrichment analysis comparing *room-temperature stored* day 0 vs. day 7 datasets. False discovery rate was corrected with Benjamini-Hochberg. "Hille-Trenk"[15] *[HT]* dataset and "Bongiovanni-Bernlochner"[16] *[BB]* dataset. Shared downregulated or upregulated genes between *HT* and *BB* are underlined.

| Gene Set | Enrichment Score | Normalized Enrichment Score | FDR Corrected p-value | Tag % | Gene % | Lead Genes |
|---|---|---|---|---|---|---|
| ***BB* Differential Expression Downregulated** | 0.822 | 2.240 | 0.01 | 1938/2335 | 18.37% | *MIEF1, SEPHS2,* ***YY1, REST, MAP2K3, EPOR, PLEKHO1, MKRN1, PANX1, SLC35E1*** |
| ***HT* Differential Expression Downregulated** | 0.805 | 2.236 | 0.01 | 877/1110 | 15.25% | ***YY1, REST, MAP2K3, EPOR, PLEKHO1, MKRN1, PANX1, SLC35E1****, CMIP, WDR1* |
| ***BB* Differential Expression Upregulated** | -0.545 | -1.889 | 0.03 | 756/2217 | 7.76% | ***TASOR2, CCDC18, ZC3H13,*** *DNAJC2,* ***NEMF, SCLT1,*** *KIF20B, SRFBP1, TCERG1, CEP63* |
| ***HT* Differential Expression Upregulated** | -0.461 | -1.747 | 0.09 | 418/1246 | 7.76% | ***TASOR2, CCDC18, ZC3H13, NEMF, SCLT1****, SENP6, ZBTB41, LUC7L3, NUP107, CSNK1G3* |

**Supplemental Figure 1. Principal components between *combined endogenous platelet aging* datasets and *room-temperature stored* datasets.** Principal component analysis was performed on *HT* and *BB* datasets individually and in a combined manner (*combined endogenous platelet aging* dataset) to compare with *room-temperature stored* datasets. Absolute cosine similarity of principal components is annotated and colored (**A**, **B**, and **C**). **A, B.** *HT* and *BB* datasets have significantly similar first principal components to the *combined endogenous platelet aging* dataset confirming that the first principal component (PC1) of the combined dataset is robust to batch effects. **C.** *Room-temperature stored* dataset has a significantly related PC1 to the PC1 of the *combined endogenous platelet aging* dataset.

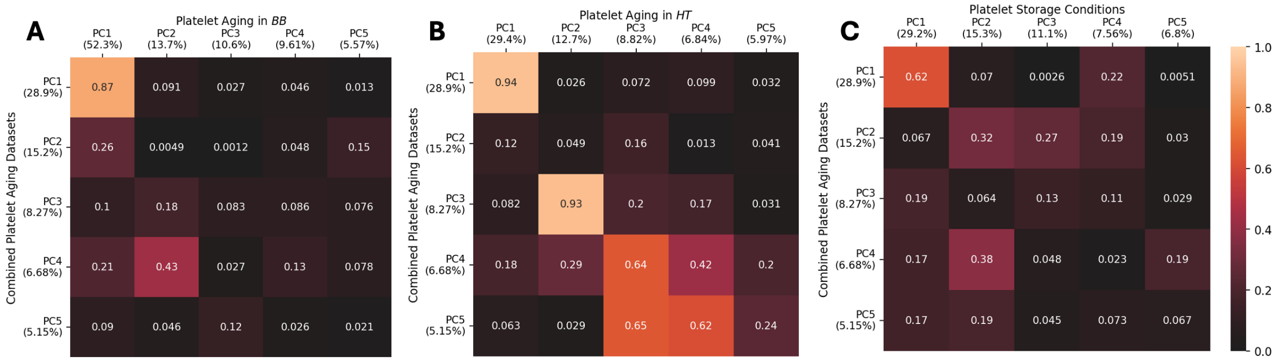

**Supplemental Figure 2. *Combined endogenous platelet aging* dataset has a consistent signature of gene alteration which is conserved in *room-temperature storage* dataset. A.** Comparison of fold change in expressed genes between *HT* and *BB* datasets and *room-temperature stored* datasets. Fold change (FC) is from young (high RNA content) platelets to old (low RNA content) platelets. In *room-temperature stored* dataset, fold change is from day 0 to day 7. Coloration is determined by direction of fold change in *HT* and *BB* datasets. Upper left, upper middle, and middle left all show fold change compared between datasets. Dashed line is a line of equivalence, not of best fit. Pearson correlation coefficient and its significance as determined by permutation are shown for each comparison. Middle, upper right, and lower right show log-2 CPM vs. log-2 fold change for expressed genes in each dataset. **B.** Comparison of fold change in DEGs between *HT* and *BB* datasets and *room-temperature stored* datasets. Plotted as in A with only genes passing differential expression in each respective comparison. **C.** When analyzing total RNA collected from platelets, an interesting problem arises: in the case of a long time-course RNA metabolism, substantial reads may be lost from one class of genes (targeted metabolism, blue) and other genes may be more inert (no metabolism, red). The final RNA total at the end of the long time-course may be much lower (e.g. 25% remaining), but the absolute read content is still seen at 100%. In this case remaining reads will fill the gap (i.e. the area under the curve must be 100% in both cases). When this occurs, reads that were in the minority previously (pie chart, left) may now be in the majority (pie chart, far right). Thus, even with no new synthesis of platelet RNA, RNA species can appear to be increasing in this context consistent with a read-displacement phenomenon.

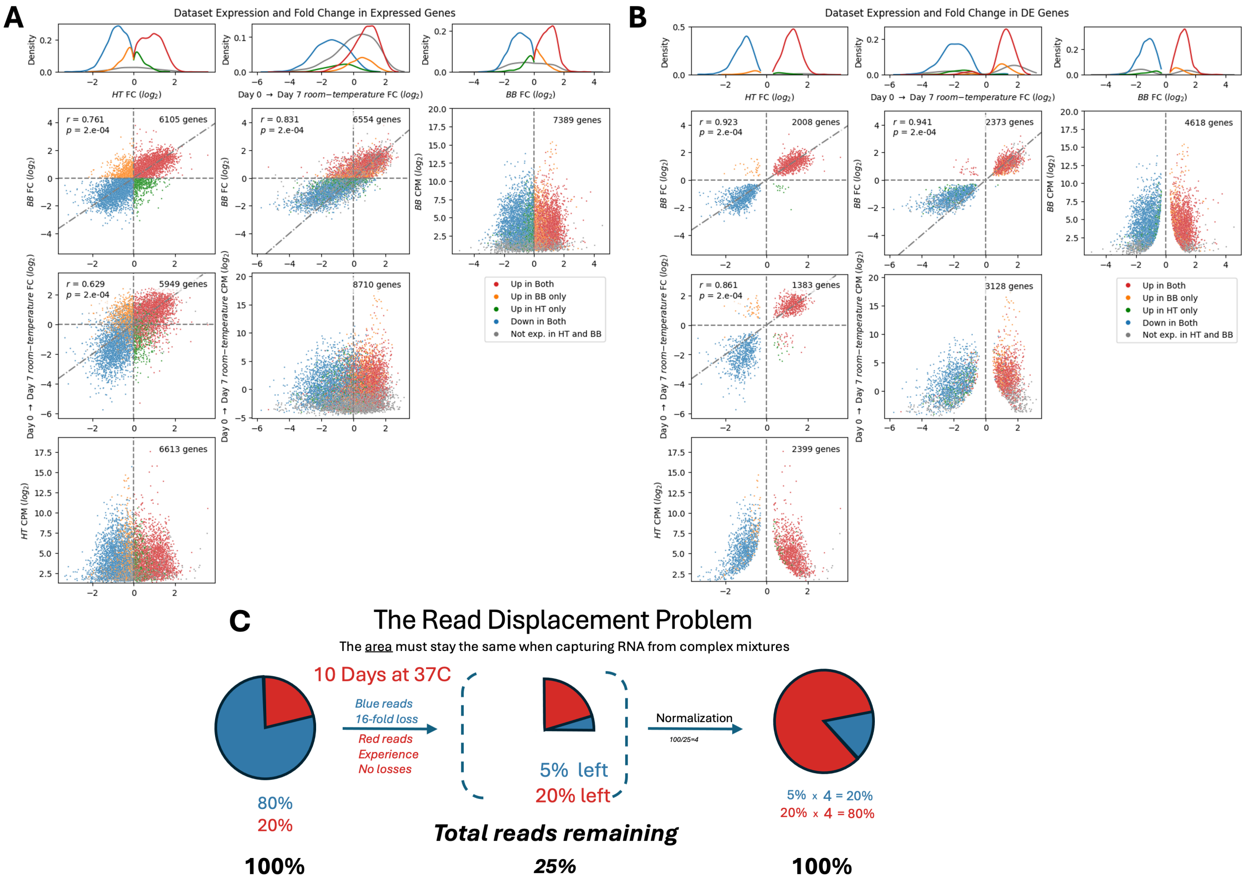

**Supplemental Figure 3. Nucleotide based counting demonstrates degradation-specific mechanism for altering gene expression in platelets**. **A.** Methods Summary: Bam alignment files are passed through a custom Python program to quantify sequenced reads at each coordinate of relevant genes. Single genes are then statistically compared between groups of samples at each point, results are tallied and displayed as a coordinate-based histogram, which we have termed "Nucleotide Level Statistical Composite" (NLSC). B-J. Condition-specific NLSC. **B**, **E**, and **H**. Top Gene Aggregated NLSC. These plots show the aggregated NLSC results for the most highly expressed genes within each condition (*BB, HT, room-temperature stored* datasets, respectively). See **Figure 2A** for differential expression details. **C**, **F**, and **I**. Positive Gene NLSC. Limited to genes with positive expression changes, these composites show genes have fewer individual points that pass statistical measures and lack the structured, 3'-directed degradation. **D**, **G**, and **J**. Negative Gene NLSC. Limited to genes with negative expression changes, these composites reveal that 3'-directed degradation is the mechanism by which those genes are decreasing in total count. **K**, **L**. Nucleotide versus Expression Changes. This shows the fraction of each gene found to change between young and old platelet positions against the fold change of overall gene expression between conditions. We determined the Spearman rank correlation coefficient of fold change and percent of gene significance and found this relationship to be significant by permutation in both *BB* (Spearman r=-0.138, p=2.5 e-3) and *HT* (Spearman r=-0.324, p=1 e-4). We see a similar 3' focused degradation pattern in both *HT* and *BB* datasets and *room-temperature stored* datasets.

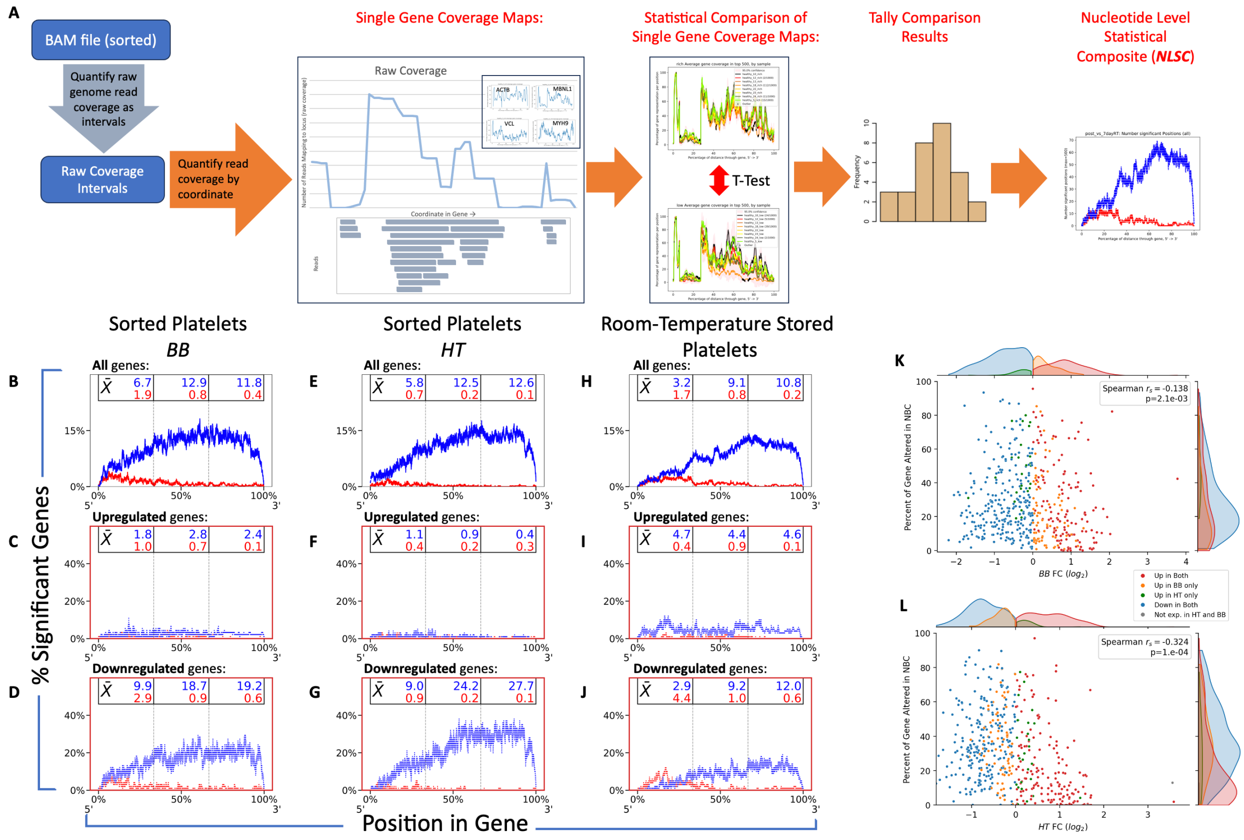

**Supplemental Figure 4. Fold change and differential expression in all *cold-stored* datasets**. **A.** Differentially expressed genes compared between young and old FACS-sorted platelet aging datasets (*HT* and *BB*), the *room-temperature* 7 day storage, and the shared DEGs of *cold-storage* timepoints compared to Day 7 *room-temperature* storage. **B** Fold change and expression for *room-temperature* storage over 7 days (copying Figure 7C for comparison). **C, F, I** Fold change comparison between FACS sorted platelet aging datasets and *cold-storage* timepoints compared to Day 7 *room-temperature* incubation (*BB* left, *HT* right; RT refers to *room-temperature*). Only genes the 669 genes shared as significantly different between *cold-storage* timepoints and Day 7 *room-temperature* storage. RT refers to *room-temperature* storage. **D, G, J** Fold change and expression of the shared 669 genes differing between *cold-storage timepoints* and *room-temperature* storage. Fold change is between Day 0 and the *cold-storage* timepoint for each period (7, 14, and 21 days respectively). No genes are differentially expressed. **E, H, K** Fold change and expression of the shared 669 genes differing between *cold-storage timepoints* and *room-temperature* storage. Fold change is between the *cold-storage* timepoint for each period (7, 14, and 21 days respectively) and Day 7 of *room-temperature* storage. All genes are differentially expressed.

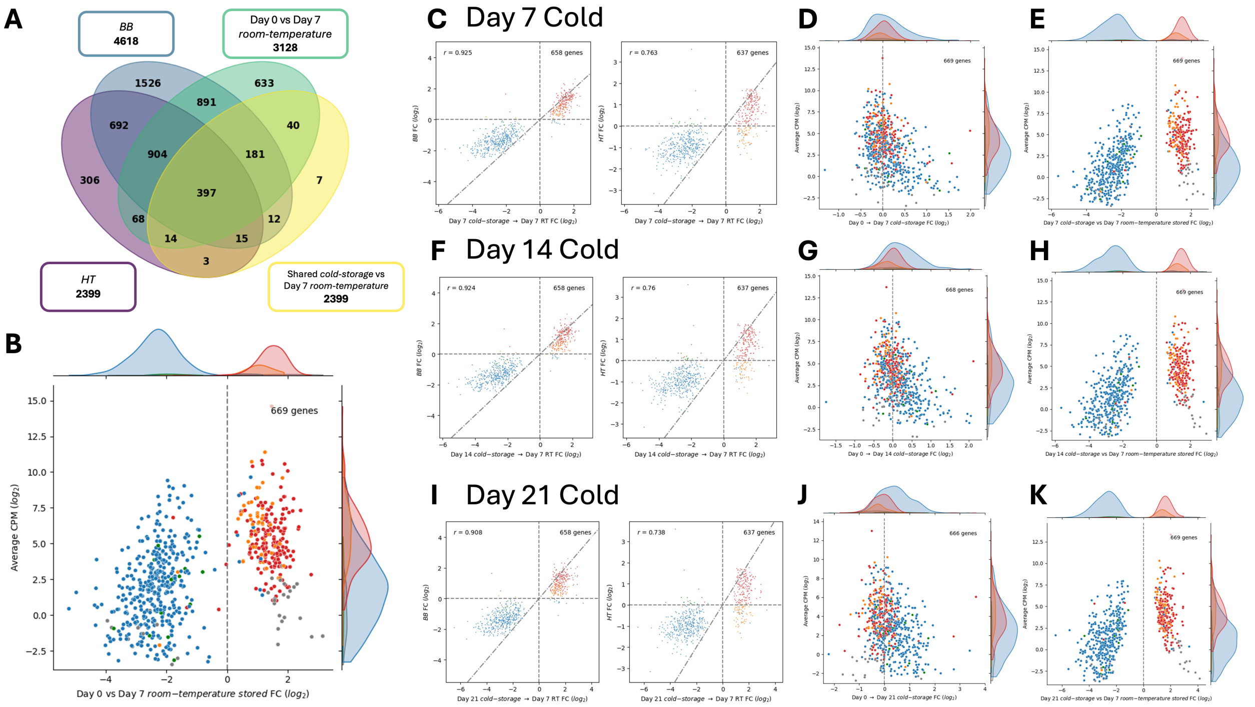


**Supplemental Figure 5. Donor variability**. **A–D**. *Individual Donor Correlations Across Storage.* Strip plots showing correlation of four alternative versions of the platelet molecular maturation scale for individuals (n=5; labelled P1, P2, etc.) across various storage conditions (room-temperature and cold-storage, for a length of time from 0 to 21 days). These alternative versions differ by dataset that they were trained on: **A**, *HT* and *BB* datasets PC1, x̄ = 0.73. **B**, *BB* dataset only PC1, x̄ = 0.64. **C**, *HT* dataset only PC1, x̄ = 0.47. **D**, *Room-temperature* and *cold-stored* datasets PC1, x̄ = 0.67. See methods for physiologic metric definitions and dataset sources.

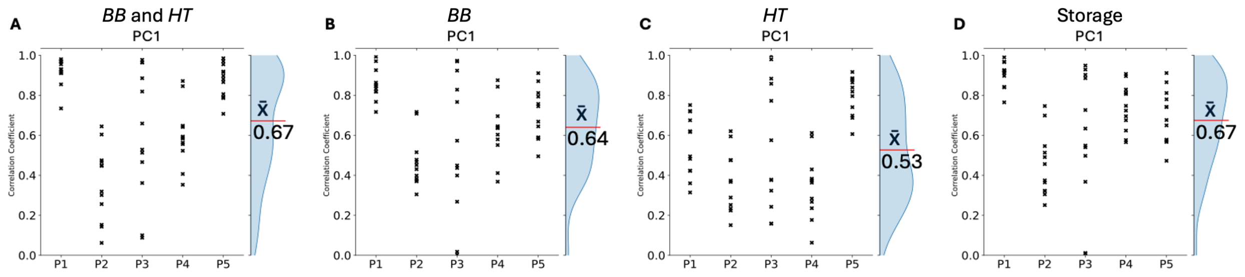

**Supplemental Table 2. Abbreviation definitions for all functional platelet measurements in Figure 5E.** Table outlines the abbreviation definitions (alphabetical by abbreviation) for all the functional measurements in **Figure 5E.**

| Abbreviation | Definition |
|---|---|
| ac | Adenosine diphosphate-collagen stimulated |
| adp | Adenosine diphosphate stimulated |
| ae | Adenosine diphosphate-epinephrine stimulated |
| alpha | Alpha angle |
| auc | Area under the curve |
| cat | Calibrated automatic thrombogram |
| cft | Clot formation time |
| col | Collagen stimulated |
| ct | Clotting time |
| epi | Epinephrine-stimulated |
| etp | Endogenous thrombin potential |
| ex | Extrinsic pathway assay |
| fib | Fibrinogen pathway assay |
| hct | Hematocrit |
| hgb | Hemoglobin |
| lag | Lag time |
| li60 | Lysis at 60 minutes |
| maxa | Maximum amplitude |
| mcf | Maximum clot firmness |
| peak | Peak thrombin generation |
| rbc | Red blood cell count |
| slp | Slope of curve |
| ttpeak | Time to peak |
| wbc | White blood cell count |
